# Genetic models of latent tuberculosis in mice reveal differential control by adaptive immunity

**DOI:** 10.1101/2021.02.08.430271

**Authors:** Hongwei Su, Kan Lin, Divya Tiwari, Claire Healy, Carolina Trujillo, Yao Liu, Thomas R. Ioerger, Dirk Schnappinger, Sabine Ehrt

**Affiliations:** Department of Microbiology and Immunology, Weill Cornell Medicine, New York, NY; Department of Computer Science and Engineering, Texas A&M University, College Station, TX

## Abstract

Studying latent *Mycobacterium tuberculosis* (Mtb) infection has been limited by the lack of a suitable mouse model. We discovered that transient depletion of protein biotin ligase (BPL) and thioredoxin reductase (TrxB2) results in latent infections during which Mtb cannot be detected but that relapse in a subset of mice. The immune requirements for Mtb control during latency, and the frequency of relapse, were strikingly different depending on how latency was established. TrxB2 depletion resulted in a latent infection that required adaptive immunity for control and reactivated with high frequency, whereas latent infection after BPL depletion was independent of adaptive immunity and rarely reactivated. We identified immune signatures of T cells indicative of relapse and demonstrated that BCG vaccination failed to protect mice from TB relapse. These reproducible genetic latency models allow investigating the host immunological determinants that control the latent state and offer opportunities to evaluate therapeutic strategies in settings that mimic aspects of latency and TB relapse in humans.

## Introduction

Infection with *Mycobacterium tuberculosis* (Mtb) most frequently results in latent tuberculosis infection (LTBI), a clinically asymptomatic state in which the only sign of infection is a positive test for immune reactivity to TB antigens, either a tuberculin skin test (TST) or interferon gamma release assay (IGRA). Individuals with LTBI are at risk for developing active TB, and many active TB cases that occur in someone with LTBI, especially in areas of low active TB prevalence, arise from reactivation of the Mtb strain that originally infected the individual. Although most active TB cases may represent progression of a primary Mtb infection that occurred recently, within 1-5 years (Behr et al., 2018, 2019), even these cases proceed through a paucibacillary, clinically asymptomatic state and therefore fit within the definition of LTBI.

TB chemotherapy often fails to sterilize Mtb infections, resulting in individuals at risk of relapse TB, even in the absence of bacterial drug resistance (Malherbe et al., 2016). In fact, 5% of TB patients that have been treated with standard TB chemotherapy develop recurrent TB (Romanowski et al., 2019; Colangeli et al., 2018; Merle et al., 2014) and recurrent TB may account for up to 10-30 % of all TB cases (Chaisson and Churchyard, 2010). In patients with recurrent TB, the Mtb strain that caused the primary infection regrows to lead to a second episode of active TB (relapse), or infection with a new strain causes the second episode (reinfection)(McIvor et al., 2017). The proportion of relapse and reinfection among recurrent TB cases varies with the overall TB incidence rate and HIV status (Cardona, 2016). Notwithstanding, post-treatment persistent infections, in which drug susceptible Mtb survives despite chemotherapy and can cause disease relapse, contributing to the spread of TB.

The standard TB mouse model exhibits some features of human TB and has informed about TB immunopathogenesis, the role of host genetics, the efficacy of antimicrobial therapy, and host-pathogen interactions (Flynn, 2006). However, it fails to mimic human LTBI, and persistent infections following TB chemotherapy. Infection of C57BL/6 or BALB/C mice by aerosol with ∼100 colony forming units (CFU) of Mtb follows a reproducible disease course consisting of a three to four week-long period during which the bacteria replicate until Mtb antigen-specific T cell responses prevent replication and maintain the infection at a stable bacterial load for many months, until eventual death of the animals (Wolf et al., 2008; Urdahl et al., 2011). During this chronic, persistent infection the typical mycobacterial load is ∼10^6^ CFU in the lung, consisting of bacilli that likely exist in different subpopulations, which are either killed by the immune system, replicating slowly, or maintain viability in a nonreplicating state (Gill et al., 2009; Wang et al., 2019). The overall bacterial burden remains nearly constant, indicating a balance between the host immune response restricting the infection and Mtb resisting eradication. Although during LTBI in humans, the bacilli are likely in a similar standoff with the immune system, a healthy human immune system restricts bacterial replication much more vigorously and is therefore capable of driving Mtb into a state of latency in which Mtb is believed to be contained at low numbers within granulomas (Barry et al., 2009; Colangeli et al., 2020). In contrast to the high-titer persistence that occurs during the chronic phase of the conventional mouse model, latent infections in humans are defined by paucibacillary stages, which are characterized by low, often undetectable numbers of Mtb and the corresponding lack of clinical symptoms. This is supported by studies in which pairs of guinea pigs were inoculated with lung and lymph node lesion tissues from people without TB symptoms, and a significant number of lesions caused infection in only one of two guinea pigs, implying a low viable burden (Opie and Aronson, 1928). Non-human primates with latent Mtb infection contained significantly fewer CFU in the lungs than animals with active disease (Lin et al., 2013). However, even animals with active disease displayed significant lesion heterogeneity.

Recently, infection of mice with an ultra-low dose of Mtb was shown to cause outcomes that more closely model human TB (Plumlee et al., 2021). Infection with 1-3 CFU resulted in heterogeneous bacterial burdens in lungs, lymph nodes and spleens and mice established pulmonary lesions that shared features with human granulomas. Some ultra-low dose infected mice developed a single granuloma with little or no dissemination to the contralateral lung. Despite the variability in bacterial burden, in all infected mice the bacterial titers increased steadily in lungs and on day 83 post infection, Mtb had disseminated to the spleens reaching titers of approximately 10,000 CFU. Thus, while ultra-low-dose infection of mice models aspects of human TB, it does not mimic relapse TB.

We sought to develop a mouse model that is characterized by low numbers of persisting bacteria and allows studying post-treatment latent infection. That Mtb can establish a latent infection in mice was first demonstrated in the “Cornell Model” developed by McCune and colleagues at Cornell University (McCune and TOMPSETT, 1956; McCune et al., 1956, 1966a; b). In this model, mice infected intravenously with Mtb were treated with isoniazid (INH) and pyrazinamide (PZA) until no bacteria could be detected by microscopy and culture on agar plates, even when homogenates of whole lungs, spleens, and livers and every organ including bones from several mice were cultured. After cessation of drug treatment, the infection relapsed spontaneously in about 30% of animals, and relapse could be induced in the remaining mice by immunosuppression with corticosteroids. The Cornell model has been used to study various aspects of Mtb persistence and latency (Flynn et al., 1998; Pinxteren et al., 2000; Dhillon and Mitchison, 1994). However, this model is notorious for its varying relapse rates and it did not result in reproducible relapse when we tried to implement it. To overcome its limitations, particularly the lack of reproducibility but also the need for prolonged antibiotic therapy which can induce resistance conferring mutations or drug tolerance (Scanga et al., 1999), we developed models in which latent infection is established via transient genetic inactivation of essential Mtb genes. After Mtb can no longer be detected, target gene expression is restored and the timing and frequency of spontaneous or corticosteroid-induced relapse report the number of mice in which Mtb persisted in the form of a paucibacillary infection. In these mouse models, Mtb persists for months below the limit of detection and reproducibly causes TB relapse in a fraction of infected animals, mimicking aspects of post-treatment latency in humans. Unexpectedly we discovered that the immunological requirements for control of latent Mtb in these models differ depending on the protein that was transiently depleted to induce latent infection.

## Results and Discussion

### Chemotherapy of Mtb infected mice did not result in consistent latent infection

In an effort to implement the Cornell model we infected C57BL/6 mice with virulent Mtb, strain H37Rv. Starting four weeks post infection, we treated the infected mice with INH and PZA for 14 weeks (Fig. S1 A). CFU in lungs, spleens, livers, lymph nodes, bone marrow and kidneys in groups of 5 mice declined with drug treatment and on day 126, when no CFU were detected in 10 mice, chemotherapy was terminated. On day 182 post infection, 8 weeks after the end of chemotherapy, a group of 10 mice was analyzed, and no Mtb was detectable in any of the mouse lungs and other organs. A group of 10 mice received dexamethasone for 4 weeks. Eight weeks later these 10 immunosuppressed mice and 10 mice that did not receive dexamethasone were analyzed for CFU, but all mice had been sterilized by the drug treatment. In a repeat experiment we increased the number of mice in each treatment group. But again, in none of the organs from 15 mice that did not receive dexamethasone was any Mtb CFU detectable, and only one out of 14 mice that had received dexamethasone relapsed with CFU in lung and spleen (Fig. S1 B). In past studies the frequency of relapse also varied substantially and sometimes relapse could not be observed at all, even with immunosuppression (Scanga et al., 1999; Lenaerts et al., 2004; Jeon et al., 2012). Therefore, we sought a different approach to reproducibly establish latent Mtb infection in mice.

### Genetic depletion of essential proteins reproducibly results LTBI in mice

We have previously developed genetic switches to control gene expression in Mtb during mouse infection and demonstrated that the depletion of essential proteins kills Mtb (Ehrt et al., 2005; Kim et al., 2013; Puckett et al., 2017; Lin et al., 2016; Tiwari et al., 2018). A dual-control (DUC) switch that combines transcriptional repression with inducible proteolysis can generate more than 99.9% repression, quickly inactivates gene activities in replicating and nonreplicating mycobacteria, and provides increased robustness against phenotypic reversion (Kim et al., 2013). We hypothesized that killing of Mtb following transient depletion of essential proteins does not fully sterilize the infection but may lead to paucibacillary infection states during which low numbers of viable bacteria persist and could reactivate to cause TB relapse. We further expected that the depletion of different targets will have different consequences for establishing a latent state and result in reproducibly distinct relapse rates. We selected DUC mutants for biotin protein ligase (BPL) and thioredoxin reductase (TrxB2) to test these hypotheses.

BPL covalently attaches biotin to acyl-CoA carboxylases and inactivation of BPL inhibits all lipid biosynthesis (Tiwari et al., 2018). Inactivating BPL is thus expected to cause pleiotropic effects, including profound changes in Mtb’s envelope structure, which might make the pathogen exceptionally vulnerable to host immunity and only rarely allow Mtb to establish a persistent, paucibacillary infection. To test this, we infected C57BL/6 mice with a BPL-DUC mutant and after the mutant had established a chronic infection, eradicated it via doxycycline (doxy) mediated BPL depletion (Fig. 1 A). Following apparent sterilization, doxycycline was withdrawn to restore BPL expression and reestablish a wild type paucibacillary infection. At multiple time points post doxy treatment groups of 10 mice were sacrificed and analyzed for Mtb CFU. We could not detect Mtb CFU in lungs or spleens of the majority of mice. In three experiments the frequency of relapse varied from 0% to 10% (mean = 6.25%). In two of these experiments, we treated additional groups of mice with dexamethasone (dexa) for 8 or 4 weeks. Eleven mice out of 35 mice that were treated with dexa relapsed; thus, the frequency of mice with regrowth in lung and/or spleen increased ∼5-fold from an average of 6.25% to 31.4%. Table 1 summarizes relapse data from multiple experiments with BPL-DUC and reveals that targeting Mtb’s BPL results in a low-frequency-relapse latency mouse model, in which almost 70% of relapse depends on immunosuppression. On average, treatment with dexa increased the relapse rate in mice infected with BPL-DUC by 25%. A Bayesian analysis of these data (see Methods) defines the 95% HDI (highest probability density interval) for the incremental effect of dexamethasone on relapse as 9.5 to 40.5%, which is statistically significant.

**Table 1.** Data obtained from 4 experiments with BPL-DUC and 6 experiments with TrxB2-DUC, which included a dexamethasone treatment group in 2 experiments with each mutant. Reported are the number of mice with regrowth in lung, in spleen or either organ, and the relapse rates.

|  | Spontaneous regrowth |  |  |  |  |  | Dexamethasone-induced regrowth |  |  |  |  |  |
| --- | --- | --- | --- | --- | --- | --- | --- | --- | --- | --- | --- | --- |
|  | number of experiments | total mice | relapsed (CFU in lung) | relapsed (CFU in spleen) | relapsed (CFU in lung or spleen) | relapse rate | number of experiments | total mice | relapsed (CFU in lung) | relapsed (CFU in spleen) | relapsed (CFU in lung or spleen) | relapse rate |
| BPL-DUC | 3 | 80 | 2 | 4 | 5 | 6.25% | 2 | 35 | 9 | 5 | 11 | 31.4% |
| TrxB2-DUC | 4 | 80 | 60 | 38 | 60 | 75.0% | 2 | 36 | 30 | 20 | 30 | 83.3% |

**Figure 1.**
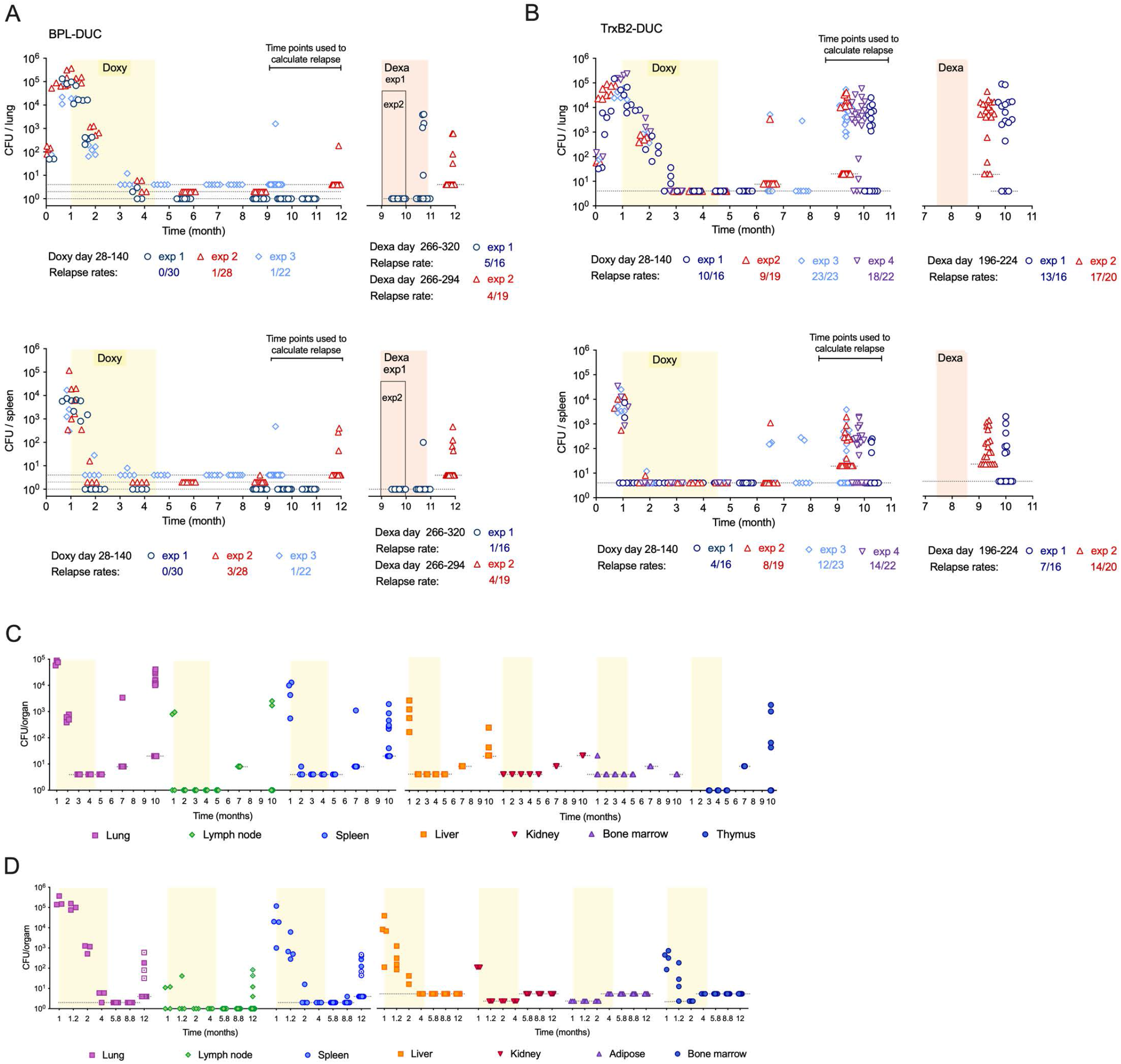
Genetic depletion of essential proteins results in reproducible paucibacillary infection. Colony forming units (CFU) in lungs and spleens of mice infected with **(A)** BPL-DUC and **(B)** TrxB2-DUC from 3 and 4 independent experiments. Mice were given doxy-containing chow (yellow area) until the infection had cleared (no detectable CFU). After that, mice were kept on regular chow. Two experiments included groups of 16-26 randomly selected mice that received dexamethasone for either the last 8 weeks or for 4 weeks starting at 2 months before the end of the experiment (pink area). Stippled lines indicate the limit of detection. **(C)** Analysis of TrxB2-DUC in multiple organs. CFU data in organs from 5 mice (months 1, 2, 3.25, 4,5) and 10 mice (month 7) and 19 mice (month 10). **(D)** Analysis of BPL-DUC in multiple organs. CFU data in organs from 4 mice (months 1, 1.2, 2, 4) and 10 mice (months 5.8, 8.8) and 38 mice (month 12), half of which were randomly selected to receive dexamethasone for 1 month starting at month 9 post infection. Data points from animals that received dexamethasone are marked by a dot within the symbol. The yellow area indicates time of doxy treatment. Stippled lines indicate the limit of detection.

Thioredoxin reductase (TrxB2) is essential for thiol redox homeostasis and has been considered as a drug target (Lin et al., 2016; Harbut et al., 2015). Depletion of TrxB2 has pleiotropic effects and kills Mtb *in vitro* and during acute and chronic high titer mouse infections. However, the importance of redox homeostasis for establishing paucibacillary infection was unclear. We analyzed the consequences of targeting TrxB2 using the TrxB2-DUC mutant in six independent experiments (Figure 1B shows data from four experiments). The experimental design was similar to the one used to analyze BPL-DUC. But in contrast to BPL-DUC, which caused on average 6.25% relapse without immunosuppression, TrxB2-DUC regrew on average in lungs and/or spleens of 75% of immunocompetent mice after a period of at least 8 weeks during which no CFU could be recovered from lungs or spleens in the majority of mice. In two experiments we included dexa treatment which caused relapse in 83.3% of the mice (Fig. 1B, Table 1). Targeting Mtb’s TrxB2 thus results in a high-frequency-relapse (>80% relapse) latency mouse model in which relapse occurs frequently even without immunosuppression. On average, treatment with dexa increased the relapse rate of mice infected with TrxB2-DUC by 10%. A Bayesian analysis defines the 95% HDI for the incremental effect of dexa on relapse for the TrxB2-DUC infected mice as -8.1 to 23.0%, which is not statistically significant.

In order to simultaneously compare the effect of strain and dexa treatment, we analyzed the combined data using a Generalized Linear Model (GLM) with a Negative Binomial likelihood function. The model included independent terms for strain and dexa treatment to predict relapse rates, along with an interaction term between them. We found, via a Wald test on coefficients, a significant difference between the reactivation rates for the two mutants (TrxB2-DUC being ∼79% and BPL-DUC only ∼19%, when averaged over dexa treatments). Furthermore, while a generalized effect of dexa on relapse rates was not significant (p=0.36) when averaged over the two strains, the model did indicate a significant interaction between dexa and BPL-DUC (p=0.0033 for the interaction term), supporting that treatment with dexa significantly increased relapse rates of mice infected with BPL-DUC (consistent with the results of the Bayesian analysis above).

For both the TrxB2-DUC and the BPL-DUC model, we confirmed that in the bacteria that were recovered following relapse, the targeted genes were still regulated in an anhydrotetracycline (atc)/doxy dependent manner and thus did not represent escape mutants. Both mutants were killed with similar kinetics in mice during doxy treatment and we confirmed that the minimal inhibitory concentration (MIC) of atc and doxy required to suppress growth in vitro was not significantly different for the two mutants (Fig. S2). Most likely, BPL depletion, which interferes with cell envelope biosynthesis synergizes more effectively with host immunity in eliminating Mtb, than depletion of TrxB2 which perturbs growth-essential pathways, including sulfur metabolism, DNA replication and antioxidant defense systems (Lin et al., 2016).

To investigate if viable Mtb was detectable in organs other than lung and spleen, we analyzed multiple organs and found Mtb in the lymph nodes, liver, kidney, bone marrow and thymus during acute infection (Fig. 1 C and D). These bacteria were eliminated with doxy treatment similarly to those in lungs and spleens and we did not detect Mtb in any of the surveyed organs during latency. We previously demonstrated that depletion of BPL and TrxB2 is accompanied by progressive healing of lesions in the lungs of infected mice (Tiwari et al., 2018; Lin et al., 2016) Previous work also demonstrated that doxy penetrates every mouse tissue and organ to the extent needed to silence Mtb genes controlled by the DUC system (Gengenbacher et al., 2020). Thus, it is unlikely that the bacilli survived in a doxy protected site.

We found that Mtb reactivated most frequently in lungs, followed by spleens and lymph nodes and thymus (Fig. 1 C and D), suggesting that the primary site of paucibacillary persistence in aerosol infected mice is the lung. We cannot distinguish whether TB relapse is caused by very few surviving organisms that may be below the limit of detection by culture on agar plates, or is due to a larger population of bacilli that are in a metabolically altered state that does not support colony formation on agar plates. We tested if Mtb was detectable with a liquid resuscitation assay (Fig. S3 A and B), that resuscitates Mtb from sputa from TB patients, chronically infected mice and *in vitro* stressed cultures (Chengalroyen et al., 2016; Hu et al., 2015; Saito et al., 2017; Imperial et al., 2018). However, this method also failed to identify viable bacilli during latency; and during active infection it yielded similar numbers as obtained by CFU determinations of infected organ homogenates on agar plates. In summary, the data presented here demonstrate that conditional depletion of an essential Mtb protein can induce latency of Mtb in mice, which is defined by a period of time lasting from weeks to months during which Mtb CFU are not detectable by culturing organ homogenates and that is followed by spontaneous or immune suppression induced regrowth of Mtb in a significant proportion of mice.

### Control of TrxB2-DUC requires adaptive immunity

These new models of TB latency provided an opportunity to interrogate the immune factors that control the Mtb latent state. T cells are crucial for the control of chronic Mtb infections (Jasenosky et al., 2015) and we therefore analyzed the pulmonary T cell populations in BPL-DUC and TrxB2-DUC infected mice at different time points of relapse experiments. As expected, the proportion of CD4 and to some degree CD8 T cells in the lung increased during acute infection and decreased with the decline of bacterial titers (Fig. 2, A-C). This included antigen 85 specific CD4 T cells (Fig. 2 D). There were no significant differences in the proportions of cytokine producing T cells between mice infected with either mutant (Fig. S3 C). We interrogated whether lung infiltrating T cells resembled effector memory T cells (T_EM_, CD44^hi^CD62L^lo^) and resident memory T cells (T_RM_, CD44^hi^CD62L^lo^CD69^+^) cells. This revealed that latent infection with TrxB2-DUC was accompanied by an increased fraction of CD4 lung-resident memory T cells and CD4 effector memory T cells compared to mice infected with BPL-DUC at five and seven months post infection (Fig. 2). Memory CD8 T cell populations were not significantly different between mice infected with the two mutants, except that at seven months post infection, the proportion of CD8 lung-resident memory T cells was higher in TrxB2-DUC infected mice compared to BPL-DUC infected mice. The increased percentage of CD4 memory T cells might reflect a larger latent TrxB2-DUC population that eventually causes relapse, than the persisting BPL-DUC population, which the mice are able to control better.

**Figure 2.**
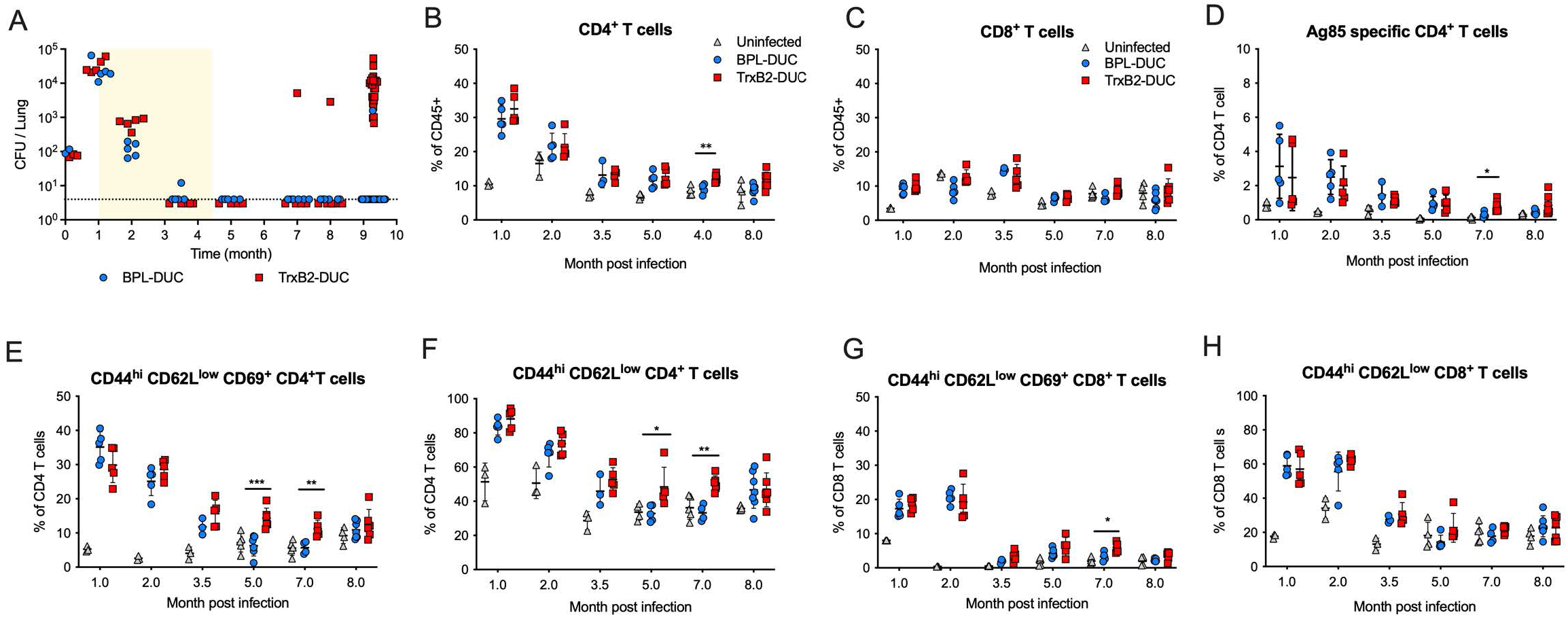
Paucibacillary infection correlates with an increased frequency of memory T cells. **(A)** CFU from lungs of mice infected with TrxB2-DUC and BPL-DUC. Data correspond to experiment 3 shown in Figure 1. The two TrxB2-infected mice that had relapsed at month 7 and 8 post infection were excluded from the analysis of T cell responses shown in panels B-H. **(B, C)** Frequency of CD4^+^ and CD8^+^ T cells in at the indicated time points. **(D)** Proportion of Ag85 specific CD4^+^ T cells. **(E, F)** CD4^+^ lung-resident memory T cells and effector memory T cells in mice infected with TrxB2-DUC and BPL-DUC. **(G, H)** CD8^+^ lung-resident memory T cells and effector memory T cells. Mice received doxycycline from month 1 to 4.5 as indicated in A. Data are from groups of 3 - 7 mice and differences in cell frequencies between TrxB2-DUC and BPL-DUC infected animals were analyzed by one-way ANOVA with Turkey’s multiple comparison test * *P* < 0.05, ** *P* < 0.01, *** *P* < 0.005.

To assess whether adaptive immunity is required to control latent Mtb, we infected SCID mice, which lack functional B and T cells, with both mutants. Because SCID mice are highly susceptible to Mtb, we started doxy treatment at two weeks post infection, kept the mice on doxy for eight weeks, which cleared the infection, and then monitored relapse (Fig. 3). BPL-DUC reactivated only in 1 of 14 mice and in this mouse could be isolated from multiple organs (Fig. 3 A and B). In contrast, TrxB2-DUC did not establish latent infection in SCID mice as it regrew almost immediately after removal of doxy and disseminated to all organs we cultured (Fig. 3 C and D). Already 8 weeks post doxy treatment, 87.5% (7 of 8) SCID mice had relapsed with CFU in their lungs (Fig. 3 C). In contrast only 13% (2 of 15) C57BL/6 mice had relapsed at 8 weeks post doxy treatment and it took roughly 20 weeks for the majority of C57BL/6 mice to relapse (Fig 1B). Thus, the majority of reactivation occurred prior to 8 weeks post doxy in SCID mice, and after 12 weeks post doxy in C57BL/6 mice. Using a Binomial statistical model, the p-value that relapse occurred at the same time in the two mouse models was 5.8e-11, strongly supporting that relapse occurred earlier in SCID than in C57BL/6 mice. This identifies the adaptive immune response as essential to establish latent Mtb infection following depletion of TrxB2. Control of BPL-DUC occurred even in the absence of adaptive immunity. The reactivation frequency of BPL-DUC in SCID mice was not significantly different from that in C57BL/6 mice (Fig. 1 A) (p=0.742 by a chi-square test), indicating that adaptive immune responses are not crucial to establish latency after inactivation of BPL. However, the differences in experimental setup, including the earlier start (2 weeks instead of 4 weeks post infection) and shorter period of doxy treatment (66 days instead of 112 days) of the SCID mice complicates the direct comparison of these experiments. Future work with identical experimental parameters is required to substantiate these findings and their interpretation. Currently, it cannot be excluded that the infrequent reactivation of BPL-DUC in SCID mice compared to TrxB2-DUC might be attributed to differences in mutant fitness or tissue distribution. Future experiments that explore the impact of innate immune cells, such as NK cells, will help shed light on the mechanisms that lead to control of this mutant in SCID mice.

**Figure 3.**
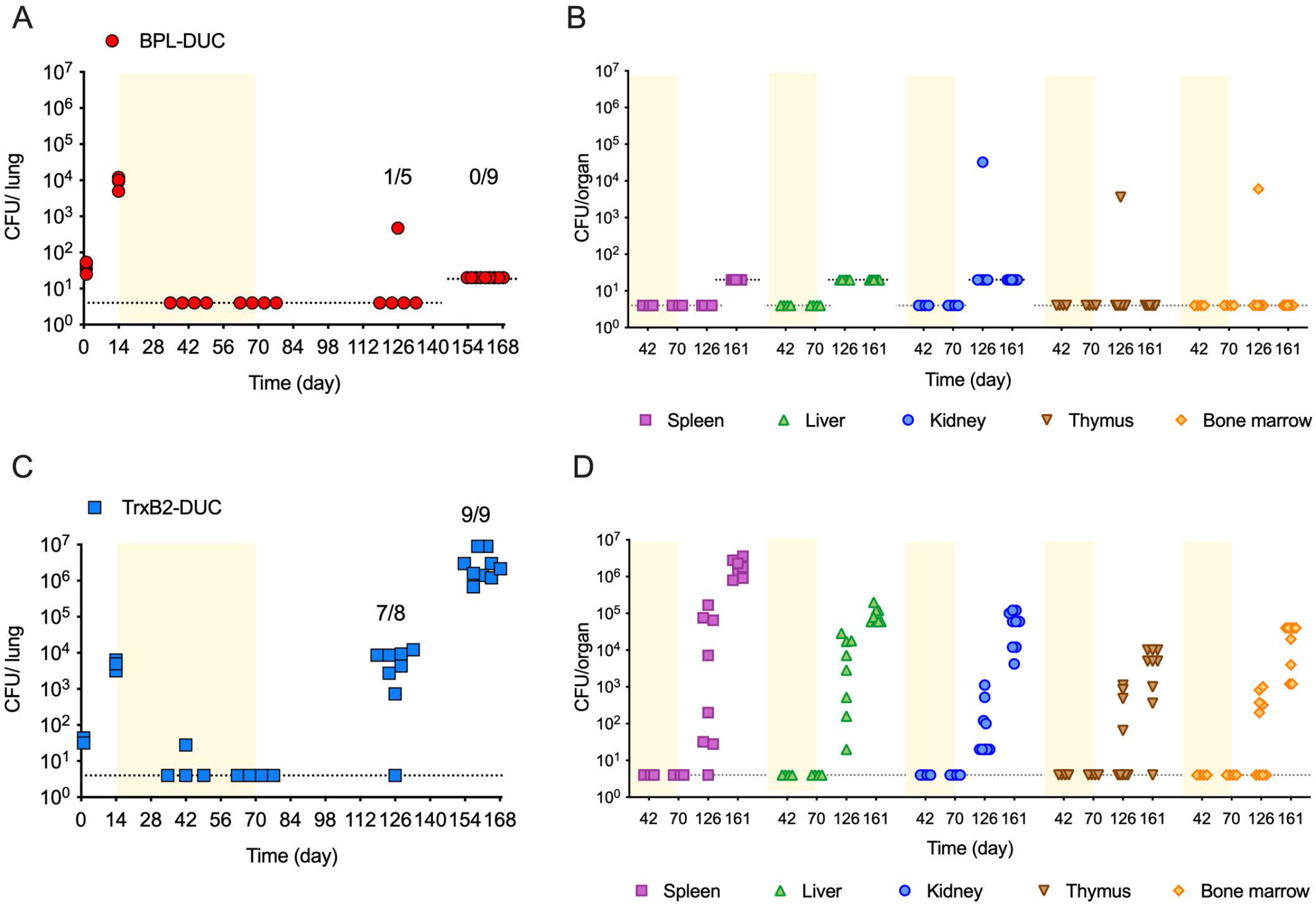
TrxB2-DUC does not establish paucibacillary infection in SCID mice. **(A)** CFU in lungs of SCID mice infected with BPL-DUC The proportion of mice with CFU is indicated. **(B)** CFU in indicated organs of SCID mice infected with BPL-DUC. One mouse relapsed with CFU in lungs and other organs on day 126. CFU in spleen and liver could not be quantified due to contamination. **(C)** CFU in lungs of SCID mice infected with TrxB2-DUC The proportion of mice with CFU is indicated **(D)** CFU in indicated organs of SCID mice infected with BPL-DUC. Mice received doxy containing food (yellow area) for 8 weeks. Stippled lines indicate the limit of detection.

### BCG vaccination does not prevent TrxB2-DUC reactivation

Vaccination with *Mycobacterium bovis* Bacille Calmette–Guérin (BCG) protects infants from extrapulmonary manifestations of Mtb infection and against progression from infection to active disease (Roy et al., 2014; Trunz et al., 2006). In contrast, its effectiveness in preventing pulmonary TB disease in adults is highly variable (Mangtani et al., 2014). We sought to determine if BCG vaccination reduces TB relapse in our genetic latency model. To assess this, we vaccinated mice that were latently infected with TrxB2-DUC with BCG Pasteur. We vaccinated via the intravenous (iv) route because in non-human primates high-dose iv BCG vaccination was recently demonstrated to prevent or substantially limit Mtb infection (Darrah et al., 2020). We analyzed immune responses in BCG vaccinated mice and phosphate-buffered saline (PBS) treated controls 25 days post vaccination and determined reactivation of Mtb in lungs and spleens of both groups on day 280 post infection, 20 weeks post vaccination (Fig. 4). BCG vaccination did not affect reactivation of latent Mtb in the lungs of mice; almost all mice in the vaccinated and the control group had CFU in their lungs (Fig. 4 A). In spleens however, Mtb regrew in only 5 of 21 vaccinated mice while 17 of 21 PBS-treated control mice yielded CFU from their spleens (Fig. 4 B). Thus, while systemic BCG vaccination did not prevent TB relapse it reduced Mtb regrowth in spleens. Vaccination might reduce dissemination of reactivating Mtb, if reactivation of Mtb primarily starts in the lung as our data suggest (Fig. 1 C and D). BCG vaccination induced more cytokine expressing CD4 T cells in the spleen than in the lung, in particular the number of IL-2 producing CD4 T cells only increased in the spleen of vaccinated mice (Fig. 4 C and D). This might be due to the higher BCG titers in spleens compared to lungs of the vaccinated mice (Fig. 4 E). BCG titers declined in the lungs approximately 10-fold from day 1 to day 24 and then remained stable until day 40 post immunization. In the spleen BCG CFU did not significantly decline during that period and remained approximately 1.5 log10 higher that in the lung. The proportion of effector memory CD4 and CD8 T cells increased in both lungs and spleens in response to BCG vaccination (Fig. 4 F and G). In summary, these data demonstrate that BCG-induced immune responses are not sufficient to prevent reactivation of latent TrxB2-DUC infection in this mouse model. The TrxB2-DUC latency model could be utilized to test vaccine candidates or strategies for their efficacy in preventing TB relapse.

**Figure 4.**
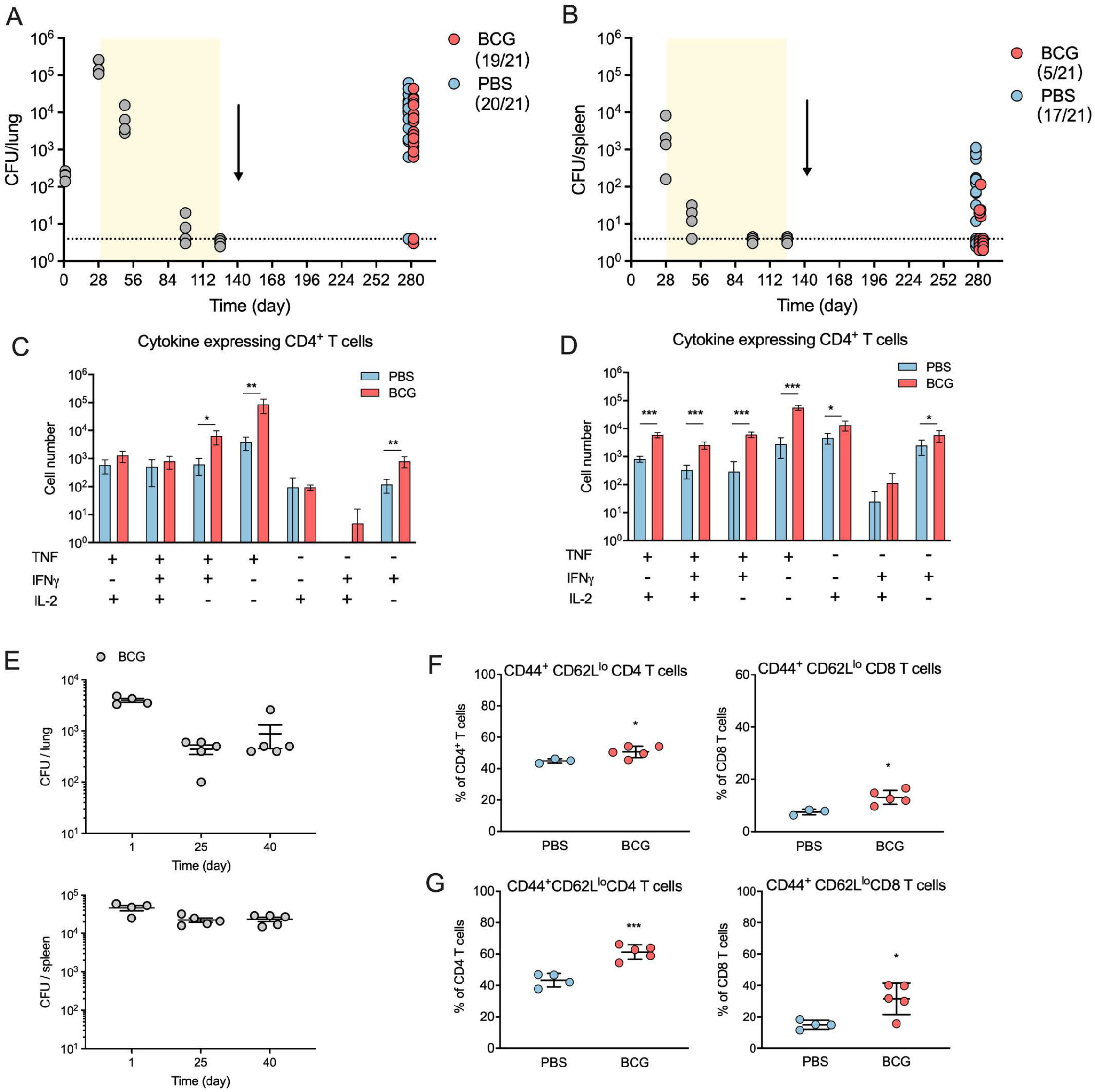
BCG vaccination fails to prevent TrxB2-DUC reactivation. **(A)** CFU from lungs and (**B**) spleens of mice infected with TrxB2-DUC and vaccinated with BCG or injected with PBS on day 140 post infection (arrow). Groups of 21 mice were analyzed on day 280 post infection, the proportion of mice with CFU is indicated in brackets. Twenty-five days post vaccination immune responses in lungs (**C**,**F**) and in spleens (**D**,**G**) were analyzed. Cytokine expressing CD4 T cells were measured by intracellular cytokine staining in CD4^+^, CD44^+^ T lymphocytes following stimulation of lung leukocytes (**C**) and splenocytes (**D**) with purified protein derivative (PPD). **(E)** Quantification of BCG CFU in lungs and spleens. On day 1, 25 and 40 post vaccination CFU in lungs and spleens were determined by culturing organ homogenates on 7H10 agar plates. (**F**,**G**) Effector memory CD4 and CD8 T cells were quantified by flow cytometry in lungs (**F**) and spleens (**G**) in groups of 3-4 PBS treated and 5 BCG vaccinated mice. The differences between PBS treated and BCG vaccinated mice were analyzed by unpaired, two tailed t-test. * *P*< 0.05; ** *P*< 0.01; *** *P*<0.001.

To address the need for mouse models of LTBI and relapse TB, we developed reproducible models of paucibacillary (low levels of bacilli) infection in mice. We demonstrated that Mtb persists for months below the limit of detection and causes relapse TB in a proportion of infected animals. Although we do not claim that these mouse models represent the equivalent of human latency, we do believe that they can inform our understanding, via comparison, of analogous human TB disease states such as reactivation from latency and relapse after TB treatment. These are both states of paucibacillary infection in which bacterial and host immune factors interact to determine whether Mtb reappears or remains clinically silent. The mouse latency models described here allow to study these aspects of TB biology that are not accessible in the conventional TB mouse model. They provide opportunity to define the host cells and molecules that prevent transition from LTBI to acute TB. They could help prioritize drug candidates that eradicate latent populations and thus could shorten chemotherapy and they could facilitate the evaluation of TB vaccine candidates.

## Materials and Methods

### Bacterial strains and culturing conditions

*Mycobacterium tuberculosis* H37Rv was used as virulent wild type strain. BPL-DUC and TrxB2-DUC have been previously described (Tiwari et al., 2018; Lin et al., 2016). BCG Pasteur was from the American Type Culture Collection strain TMC 1011 (ATCC #35734). Strains were cultured in liquid Middlebrook 7H9 medium supplemented with 0.2% glycerol, 0.05% tween80 and ADN (0.5% bovine serum albumin, 0.2% dextrose, 0.085% NaCl) and on Middlebrook 7H10 agar plates supplemented with 0.2% glycerol and Middlebrook OADC enrichment (Becton Dickinson). Antibiotics were added for selection of genetically modified strains at the following concentrations: hygromycin (50 μg/ml), kanamycin (25 μg/ml), zeocin (25 μg/ml).

### Mouse infections

Female 8- to 10-week-old C57BL/6 (# 000664, Jackson Laboratory) or SCID mice (# 001913, Jackson Laboratory) were infected with the indicated Mtb strains using an inhalation exposure system (Glas-Col) with a mid-log phase Mtb culture to deliver approximately 200 bacilli per mouse. To vaccinate, mice were injected in the lateral tail vein with 100 μl BCG suspension in PBS at a concentration of 10^7^/ml. Mice received doxycycline containing mouse chow (2,000 ppm; Research Diets) for the indicated periods. For the Cornell model, mice received INH (125 μg/ml) and PZA (15 g/L) in the drinking water for 14 weeks. To enumerate CFU, organs were homogenized in PBS and cultured on 7H10 agar. Charcoal (0.4 %, w/v) was added to the plates that were used to culture homogenates from doxy and antibiotic treated mice. Agar plates were incubated for 3-4 weeks at 37°C. We identified suppressor mutants by culturing half of each organ homogenate on 7H10 plates containing atc (500 ng/ml). When suppressor mutants grew up, which was very rare, the infected mice were excluded from the experiment. Mice were housed in a BSL3 vivarium. All mouse experiments were approved by and performed in accordance with requirements of the Weill Cornell Medicine Institutional Animal Care and Use Committee.

### Limited dilution assay and MPN calculations

Half of each lung homogenate (3 right lung lobes resuspended in 2 mL PBS supplemented with 2% (v/v) BD MGIT PANTA antibiotic mixture, BD Cat#245114) was used to set up the limited dilution series as described previously (Saito et al., 2017). Mtb H37Rv culture filtrate was prepared and MPN calculations were performed as described (Saito et al., 2017).

### Flow cytometry

Mouse lungs were isolated and placed in RPMI1640 containing Liberase Blendzyme 3 (70 μg/ml; Roche) and DNase I (50 μg/ml; Sigma-Aldrich). Lungs were then cut into small pieces and incubated at 37°C for 1 hour. The cells were filtered using cell strainers, collected by centrifugation, resuspended in ACK hemolysis buffer (ThermoFisher) and incubated for 10 minutes at room temperature. Cell were then washed with PBS and resuspended in splenocyte medium (RPMI-1640, supplemented with 10% FBS, 2 mM GlutaMax 10 mM HEPES, and 50 μM M2-mercapto-ethanol). For intracellular staining samples, cells were stimulated with PPD (20 μg/ml) in the presence of anti-CD28 antibody (37.51, BioLegend) for 1.5 hours and 10 μg/ml Brefeldin A (Sigma) and monesin were added and incubated at 37 degree for another 3 hrs. Samples were kept on ice in a refrigerator overnight. Cells were stained with Zombie-UV (BioLegend) to discriminate live and dead cells. Purified anti-CD16/32 antibody (93, BioLegend) was used to block Fc receptor before staining. PerCp-Cy5.5 anti-CD62L (MEL-14, Thermofisher), Alexa 700 anti-CD45 (30-F11, BioLegend), BV605 anti-CD4 (RM4-5, BioLegend), BV711 anti-CD8 (53-6.7, BioLegend), APC Cy7 anti-CD69 (H1.2F3, BioLegend), BUV395 anti-CD44 (IM7, BD Biosciences) were used to stain cells for 30 minutes at room temperature. PE-conjugated I-A(b) Mtb Ag85B precursor 280-294 (FQDAYNAAGGHNAVF) from the National Institutes of Health tetramer core facility, Atlanta, GA were used to stain for Ag85 specific T cells. The cells were fixed in fixation buffer (BioLegend) for 30 minutes and taken out of BSL3. The cells were incubated for 20 minutes in permeabilization buffer (eBioscience) before intracellular cytokine staining. FITC anti-TNF (MP6-XT22, BioLegend), BUV737 anti-IFNγ (XMG1.2, BD Biosciences), APC anti-IL-2 (JES6-5H4, BioLegend), BV785 anti-CD3 (17A2, BioLegend) were used to stain cells for 30 minutes. Cells were washed and resuspend in cell staining buffer (BioLegend). Flow cytometry data were acquired on a cytometer (LSR Fortessa TM; BD Biosciences) and were analyzed with FlowJoTM V10. Gating strategies are depicted in Fig. S3.

### Statistical Analysis

Flow cytometry data were analyzed by one-way ANOVA with Turkey’s multiple comparison test or by unpaired, two tailed t-test and differences in cell frequencies between TrxB2-DUC and BPL-DUC infected animals are reported using Prism 9.0.1 (GraphPad Software).

Effect of dexa on relapse rates for BPL-DUC and TrxB2-DUC: Dexa treatment increased the relapse rate for both mutants: by 25% for BPL-DUC, and around 8% for TrxB2-DUC. To assess the statistical significance of these increases (or ‘deltas’), we employed a Bayesian model to calculate a *confidence interval* for each delta (to compare to 0). We modeled the underlying relapse rate ϕ for each mutant with and without dexa using a Binomial distribution as a likelihood function with a *Beta(α,β)* conjugate prior. Therefore, the joint probability distribution of the data is:

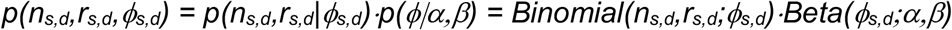

where *n*_*s,d*_, is the total number of mice infected with strain *s* and where *d* indicates dexa treatment (yes or no), and *r*_*s,d*_ is the number of mice from this arm that relapsed. It can be shown that the posterior distribution over the relapse rate ϕ is given by an updated Beta as: *p(ϕ* | *n,r) = Beta(α+r,β+n-r)*

In order to determine a posterior distribution on the deltas (dexa effect for each strain), we first generated a representative sample from the posterior of each relapse rate variable by Monte Carlo sampling (10,000 random draws from the updated Beta posterior distributions, using α=1,β=1 for the hyperparameters of the Beta prior), and then took the difference of samples to generate the distribution over the deltas: δ_s_ ∼ ϕ_s,d=yes_ - ϕ_s,d=no_

To quantify the statistical significance, the 95% HDI (highest probability density interval) was extracted from the Monte Carlo sample representing the posterior distribution for each delta. The 95% HDI for the incremental effect of dexa on reactivation of BPL-DUC is 9.5 to 40.5%, which is statistically significant (because it does not overlap 0). The 95% HDI for the incremental effect of dexa on relapse for the TrxB2-DUC infected mice is -8.1 to 23.0%, which is not statistically significant (because it overlaps 0). This analysis supports that dexamethasone treatment has a significant effect on the relapse rate for BPL-DUC but not for TrxB2-DUC.

Linear Model for Evaluating Combined Effects of Strain and Dexamethasone Treatments: To extract additional insights from the mouse data and quantify the significance of trends, we performed an analysis of the combined mouse data with a Generalized Linear Model (GLM) using a Negative Binomial likelihood function (with log link). The model has independent terms for strain and dexa treatment to predict relapse rates:

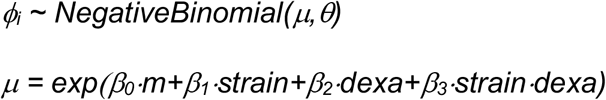

where *strain* and *dexa* are binary variables encoding the experimental conditions, and *m* is the number of mice in each experiment. The Negative Binomial has two parameters: mean μ, and shape parameter θ (inverse of dispersion). The (log of the) mean number of relapsed mice in each experiment is fit to a linear equation as a function of mice, strain, and dexa, with an interaction term between strain and dexa; the shape parameter θ is optimized internally from the data. The model was fit using *glm*.*nb*() in R.

In the resulting model, the difference between the relapse rates for the two mutants (TrxB2 being ∼79% and BPL only ∼19% when averaged over dexa treatments) was found to be weakly statistically significant (p=0.0664 by a Wald test). Furthermore, while a generalized effect of dexa on relapse rates was not significant when averaged over the two strains (p=0.36), the model *did* indicate a significant interaction between dexa and the BPL mutant (p=0.0033 for the interaction term), supporting that treatment with dexa significantly increased relapse rates of mice infected with the BPL mutant (consistent with the results of the Bayesian analysis above).

Analysis of difference in timing of relapse of TrxB2-DUC infected C57BL/6 vs SCID mice: TrxB2-DUC infected SCID mice relapsed with high frequency (∼90%, similar to the relapse rate in C57BL/6 mice at ∼75%; no dexamethasone), and relapse occurred early. Data in Figures 1b and 3c, show that the majority of SCID had relapsed by 8 weeks after doxy treatment, whereas relapse of the majority of C57BL/6 mice occurred after 12 weeks post-doxy. In order to assess the statistical significance of this difference in timing of relapse, we tested the Null hypothesis that relapse occurred in C57BL/6 mice as early as in SCID mice (<8 weeks) and that the low proportion of mice with CFU in C57BL/6 mice at weeks 8 and 12 post doxy was just due to sampling error. The maximum likelihood estimate of the longer relapse rate of C57BL/6 mice is 75.0% (60/80, based on pooling the mice relapsed at 20 and 23 weeks post doxy). Assuming reactivation had occurred by week 8 post-doxy in C57BL/6 mice, the p-value for the outcomes observed at weeks 8 and 12 post doxy (combined, 3 out 25 mice relapsed) is the cumulative of the Binomial for r≤3 given n=25 and p=0.75, which is p=5.8e-11, strongly supporting that relapse occurred later in C57BL/6 than SCID mice.

## Online supplemental material

Fig. S1 displays the data from the Cornell mouse model experiments. Fig. S2 displays data demonstrating that BPL-DUC and TrxB2-DUC are equally susceptible to anhydrotetracycline and doxycycline mediated growth inhibition. Fig. S3 shows data revealing that there is no significant difference in detecting TrxB2-DUC by CFU and liquid outgrowth assay and no differences in cytokine producing CD4+ T cells in lungs from mice infected with BPL-DUC and TrxB2 DUC. The figure also shows the gating strategies for CD4^+^ and CD8^+^ T cells and representative flow cytometry plots identifying memory T cells.

## Acknowledgements

We thank J. Andres, N. Betancourt, I. DaSilva, J. Kim, J. McConnell, R. Aguilera Olvera, N. Song, P. Pino Tamayo for technical assistance. We thank K. Saito, T. Warrier, K. Burns, C. Nathan, M. Glickman, K. Rhee and V. Dartois for advice and helpful discussions; M. Glickman, C. Nathan and K. Rhee for critically reviewing this manuscript and constructive suggestions. This work was supported by the Tri-Institutional TB Research Unit (NIH Grant U19 AI111143).

## Author contributions

H. Su, K. Lin, D. Tiwari, C. Healy, C. Trujillo, Y. Liu, D. Schnappinger, and S. Ehrt conceived the study and analyzed the data; H. Su, K. Lin, D. Tiwari, C. Healy, C. Trujillo, Y. Liu performed the experiments; T.R. Ioerger performed statistical analyses. S. Ehrt wrote the manuscript with input from the other authors.

The authors declare no competing financial interests.

## Figure Legends

**Figure S1.**
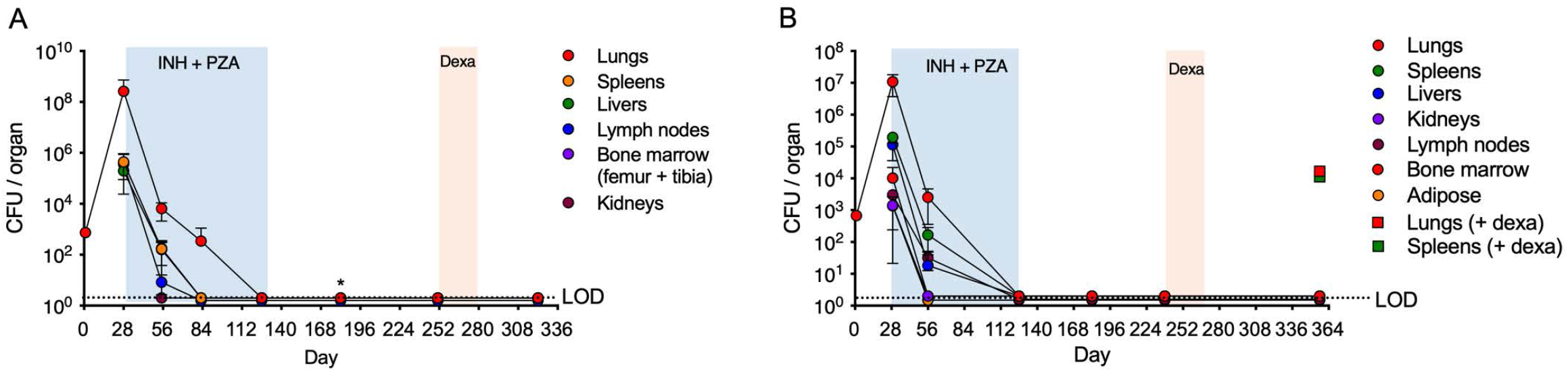
The TB Cornell Model does not result in reproducible paucibacillary infection. Colony forming units (CFU) in the indicated mouse organs of mice infected with H37Rv. Two experiments **(A, B)** were conducted. Mice received isoniazid (INH) and pyrazinamide (PZA) for 14 weeks (blue area) and a group of 10 (experiment A) and 14 (experiment B) mice received dexamethasone for 4 weeks (pink area). Before and during chemotherapy 5 mice were sacrificed at each timepoint to determine CFU in the indicated organs. At the end of and after chemotherapy 10-15 mice were analyzed in each treatment group. Stippled lines indicate the limit of detection. In experiment A, one mouse yielded one CFU in its liver homogenate on day 182 (indicated by *). In experiment B, one dexamethasone-treated mouse relapsed with CFU in lung and spleen.

**Figure S2.**
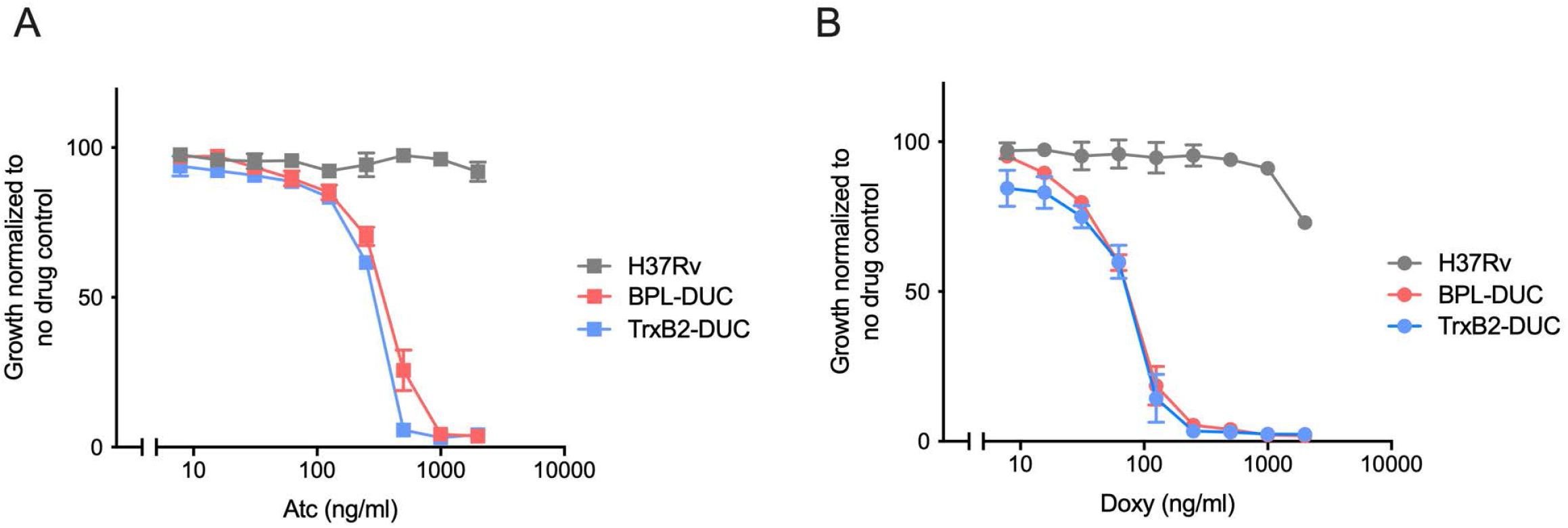
BPL-DUC and TrxB2-DUC are equally susceptible to anhydrotetracycline and doxycycline mediated growth inhibition. Impact of **(A)** anhydrotetracycline (atc) and **(B)** doxycycline (doxy) on growth of H37Rv, BPL-DUC and TrxB2-DUC. Data are means ± SD of triplicate cultures and representative of three independent experiments.

**Figure S3.**
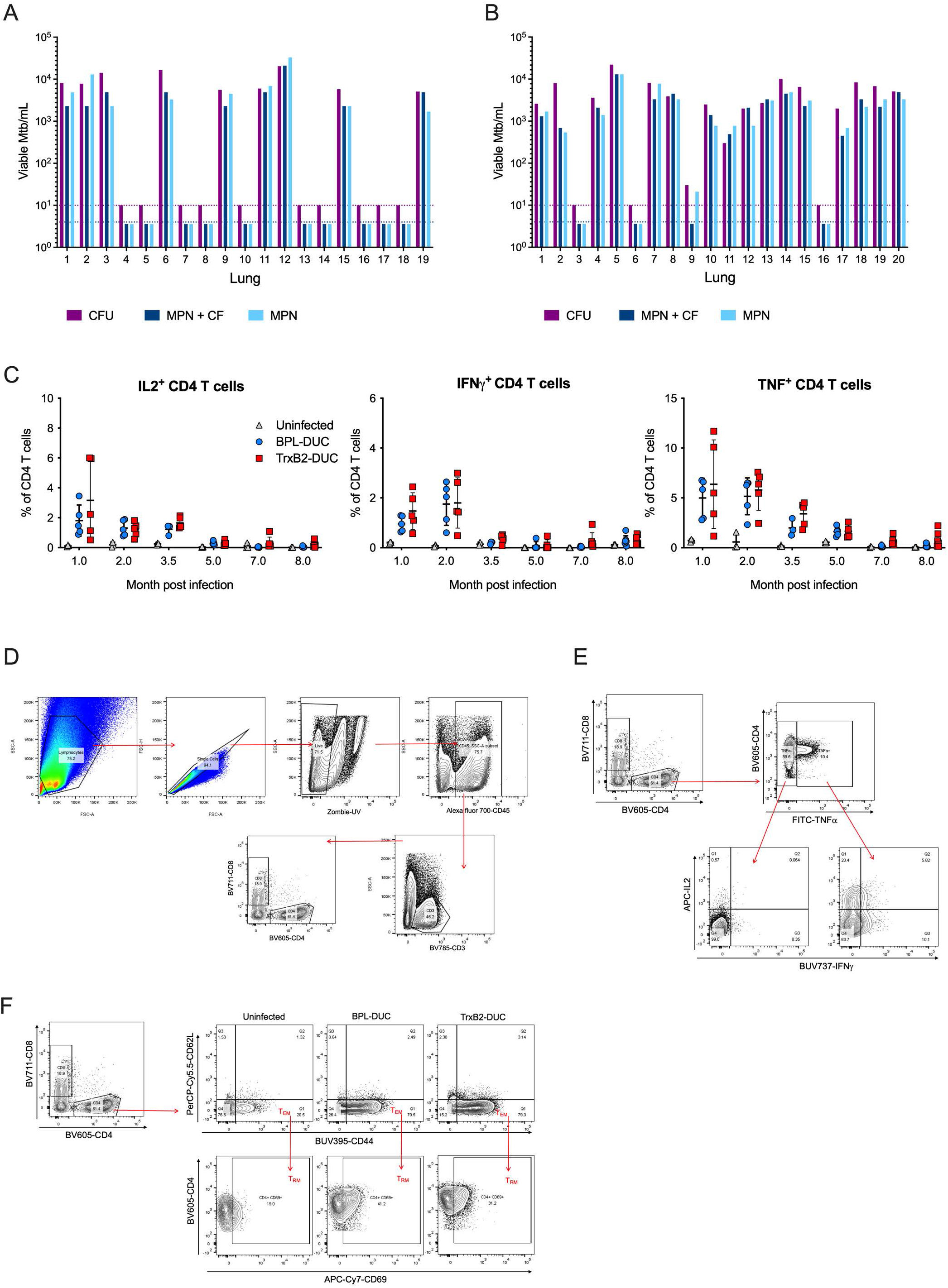
No significant difference in TrxB2-DUC detection by CFU and liquid outgrowth assay and no differences in cytokine producing CD4+ T cells in lungs from mice infected with BPL-DUC and TrxB2 DUC. **(A)** Analysis of TrxB2-DUC in mouse lungs 10 months post infection. Viable Mtb in lungs from 19 mice that did not receive dexamethasone and **(B)** in lungs from 20 mice that received dexamethasone for 4 weeks. Each lung was split in 3 parts and analyzed by conventional CFU assay or liquid outgrowth assay to estimate the “most probable number” (MPN) with or without addition of culture filtrate (CF) Stippled lines indicate the limit of detection. (**C**) Quantification of cytokine producing CD4^+^ T cells in lungs from uninfected mice and mice infected with BPL-DUC and TrxB2 DUC at the indicated time points. Percent cytokine positive T lymphocyte responses as measured by intracellular cytokine staining in CD4^+^, CD44^+^ T lymphocytes following stimulation of lung leukocytes with purified protein derivative (PPD). Mice received doxycycline from month 1 to 4.5 (see Fig. 2 A). Data are from groups of 3 -7 mice. **(D)** Gating strategy for CD4^+^ and CD8^+^ T cells in lungs infected with TrxB2-DUC at day 28. **(E)** Gating strategy for multifunctional CD4 T cells. **(F)** Representative flow cytometry plots identifying memory T cells in lungs of uninfected mice and mice infected with BPL-DUC and TrxB2-DUC on day 28 post infection.

